# The multilayered organization of water-soluble proteins

**DOI:** 10.1101/2021.07.07.451496

**Authors:** Lincong Wang

**Affiliations:** The College of Computer Science and Technology, Jilin University, Changchun, Jilin, China

**Author notes:** Corresponding author: Lincong Wang.

**Keywords:** Solvent-excluded surface, Protein interior, Multilayered organization, Surface charge, Protein-solvent interaction, Solvation

## Abstract

The structural analysis of proteins has focused primarily on secondary structure, three-dimensional fold and active site while whole surface has been analyzed to a lesser extent and interior has not received much attention. Here we present an analysis of both the surfaces and the interiors of a set of water-soluble monomeric proteins in terms of solvent-excluded surface (SES) and atomic partial charge. The analysis shows that the surface of a soluble monomer has a net negative charge and is much smoother than the interior. Most interestingly with regard to both atomic partial charge and SES-defined geometric property there exists a multilayered organization from the exterior to the interior of a soluble monomer. The multilayered organization is closely related to protein-solvent interaction and should be a general feature of a water-soluble protein. Particularly the multilayered organization may set an upper limit for the size of a water-soluble monomer and plays an important role in the determination of its overall shape in solution.

**Significance statement:** The analysis of the solvent-excluded surfaces (SESs) of a large set of water-soluble monomers with crystal structures shows that in any soluble monomer there exists a multilayered organization in terms of SES-defined electric and geometric properties. It means that the atoms in a soluble monomer are not randomly distributed but organized into successive layers to optimize its interaction with solvent molecules. The SES-derived multi-layered organization should be a general feature of a water-soluble protein and likely plays a vital role in its solvation, folding and structure.

## 1 Introduction

The experimentally-determined structures of proteins have been analyzed in depth to rationalize their physical-chemical properties such as solvation and folding and their biochemical properties such as binding affinity and enzymatic parameter. Previous analyses though extensive focus mainly on either secondary structure contents, or three-dimensional folds, or the physical-chemical and geometric properties of binding sites or active sites or protein-ligand interfaces. By comparison protein surface as a whole has not been as well studied. For example no accurate quantitative relationships have been established between the physical-chemical and geometrical properties of the surface of a protein and its solvation, folding and structure [1, 2, 3, 4, 5, 6]. At present it is challenging to evaluate the contribution of surface to solvation using either experimental approach [7, 8] or theoretical model [9, 10, 11, 12, 13] or molecular dynamic (MD)^1^ simulation [14, 15] or structural information [16, 17]. For example due in part to the difficulty to quantify protein-solvent interaction [18] it is not clear at present whether or not and how the surface of a water-soluble naturally-occurring protein has best adapted to aqueous solvent. Clues to possible adaptation may be obtained through a systematic and detailed analysis of both the surfaces and the interiors of the proteins with known structures.

There exist three models for molecular surface called van der Waals (VDW) surface, solvent accessible surface (SAS) [19, 20] and solvent-excluded surface (SES) [20, 21]. An SES is a two-dimensional (2D) manifold that is impenetrable to solvent molecules. In other words an SES determines a 2D boundary that seals off the interior of a protein from direct contact with solvent molecules. SES consists of three different types of 2D patches: convex spherical polygon on a solvent-accessible atom, saddle-shaped patch on a torus defined by two accessible atoms^2^ and concave spherical patch on a probe determined by three accessible atoms. Their surface areas are called respectively SAS area, torus area and probe area.

To examine SES’s contribution to protein solvation, folding, stability and structure, and to obtain clue to the adaptation of soluble proteins to aqueous solvent we have applied our accurate SES computation algorithm to a set 𝕄 of 4, 290 water-soluble monomers with crystal structures and a set 𝕌 of 1, 585 structure models of extended conformations. The latter is used only for reference. For each soluble monomer we compute not only the SES for the whole protein but also the SESs for successive *interior* layers^3^. For clarity the set of solvent-accessible atoms for a whole protein will be called *exterior* SES layer or exterior layer in short. An interior SES layer is defined as the set of newly-exposed atoms after the removal of the layer of surface atoms determined by the previous SES computation. For example the first interior layer of a protein is the set of exposed atoms after the removal of the exterior SES layer. With each surface atom in either exterior or interior layer we associate an atomic area defined as the summation of its SAS area, torus area and probe area. The atomic SES areas of a set of protein atoms could be used to divide the atoms into a layer of surface atoms with nonzero SES area and a corresponding core of buried atoms with zero SES area. Several electric and geometric properties likely to be relevant to protein-solvent interaction are then defined to characterize both the exterior and interior SES layers for soluble monomers and the exterior SES layers for extended conformations. As summarized below a key result of our analysis is that these SES defined-electric and geometric properties change systematically rather than randomly from exterior layer to the innermost interior layer. In other words in terms of these SES-defined properties there exists in a soluble protein from the exterior to the innermost core a multilayered organization.

With the division of a set of atoms with partial charges into a layer of surface atoms and a core of buried atoms we could analyze the distribution of atomic partial charges in a protein. The analysis shows that for every monomer in 𝕄 the partial charge per surface atom *ρ*_*s*_ for the exterior layer is *negative* and the *ρ*_*s*_s for the successive interior SES layers vary in a *zigzag* manner and with a trend towards more positive. In the contrary the charge per buried atom *ρ*_*b*_ for the core buried by the exterior layer of a soluble monomer is *positive* though the *ρ*_*b*_s for the successive cores also become more and more positive. The result that the surface of a soluble monomer has a net negative partial charge provides at atomic level a piece of evidence for the conclusion that polar and charged residues prefer to be on the surface of a water-soluble protein [22, 23, 24, 25].

To characterize the geometry of an SES layer we introduce a geometric property called concave-convex ratio *γ*_*cc*_ defined as the ratio of the summation of its torus area and probe area over its SAS area^4^. Our analysis shows that for every soluble monomer *γ*_*cc*_ decreases invariably from the exterior layer to the innermost interior layer. Furthermore the *γ*_*cc*_ for an exterior SES layer differs largely from those for interior SES layers. Similarly for every soluble monomer SES area per atom increases constantly from exterior to the interior^5^. The continuous increase in SES area per atom and the continuous decrease in *γ*_*cc*_ from exterior to the interior are consistent with the conclusion that the core of a folded protein is tightly packed [26]. On the other hand a multilayered organization with continuous increases from exterior to the innermost interior in both packing and net partial charge may set an upper limit for the size of a water-soluble monomer and by extension may also determine its overall shape in solution.

Previously it has been shown that the intermolecular VDW attractions between accessible apolar atoms and solvent molecules [27] and the intermolecular hydrogen bonds (H-bonds) between accessible polar atoms and solvent molecules play vital roles in protein solvation [28]. To analyze their contributions to protein-solvent interaction we first divide the atoms of an exterior layer into a set of apolar atoms and a set of polar atoms^6^ with the latter being further divided into a set of atoms **H**_b_ that form at least one H-bond with a protein atom and a set of atoms **H**_nb_ that form no such an H-bond. The analysis shows that there exist large differences in SES area per atom and concave-convex ratio between apolar atoms and polar atoms and between the polar atoms in **H**_nb_ and those in **H**_b_. The differences could be explained by best adaptation of a soluble protein to aqueous solvent through the optimizations of both the VDW and the hydrogen bonding (H-bonding) interactions between protein surface atoms and solvent molecules. The optimizations may in turn lead to the formation of the multilayered organization in a soluble monomer.

## 2 Materials and Methods

In this section we first describe the data sets and then briefly present SES computation. Finally we describe a list of SES-defined electric and geometric properties that are likely relevant to protein-solvent interaction.

### 2.1 The data sets

We have downloaded from the PDB a non-redundant set of crystal structures of water-soluble monomeric proteins each has a resolution ≤ 3.5Å and a *R*_free_ ≤ 27.5%. This set excludes membrane proteins and nucleic acid binding proteins in order to minimize other structural features that may contribute to protein-solvent interaction. A prepossessing step that requires no structure to have > 5% missing atoms^7^ and no structure to include a bound compound with > 20 heavy atoms selects a set 𝕄 of 4, 290 structures. The structures in 𝕄 have the number of atoms (including protons) ranging from 817 to 27, 859. The sequence similarities among these monomers are rather low (section S1 of Supporting Information (SI)). Out of 𝕄 we select a subset 𝔽 of 1, 585 structures with 1, 008 to 10, 313 atoms that have no gap in sequence^8^, < 0.2% missing atoms and no bound compounds with > 5 heavy atoms. They are used to represent soluble proteins in native state. Their corresponding unfolded states are represented by a set of extended and energy-minimized conformations 𝕌 generated by CNS [29] using the amino acid sequences in 𝔽.

### 2.2 The preprocessing of PDB files for SES computation

The downloaded PDB files are preprocessed as follows for SES computation. Protons are first added using the program REDUCE [30] to any PDB that lacks their coordinates and the protonated structures are then analyzed by our structural analysis and visualization program. A graph with atom as node and bond as edge is first constructed for each of the 20 naturally-occurring amino acid residues, Hsd, Hsp and protonated Asp and Glu residues using CHARMM atom nomenclature [31]. A molecule graph is then built for a whole protein by adding an edge for each peptide bond. For an atom with more than one conformation, only its first form is selected for SES computation. Next any gap in a protein chain is identified and the percentage of missing atoms in each structure is computed by a comparison of the atoms in the PDB file with those in the protein molecule graph. CHARMM force field parameters (e.g. partial charges) [31] are assigned to individual atoms using protein molecule graph. Only protein atoms are included in SES computation.

### 2.3 SES computation

An SES is composed of three types of areas: *a*_*s*_, *a*_*t*_, *a*_*p*_ where SAS area *a*_*s*_ is the area of an exposed spherical polygon on the surface of an accessible atom, torus area *a*_*t*_ is the area of a patch on a torus defined by two atoms and two probes, probe area *a*_*p*_ is the spherical polygon area on the surface of a probe whose position is determined by three atoms. For an isolated probe that does not intersect with any other probes the polygon reduces to a spherical triangle but for a probe intersecting with other probes the polygon may assume any shape. Both SAS area and torus area are computed analytically with the former computed using Gauss-Bonnet theorem. Probe area is also computed using Gauss-Bonnet theorem except for very rare cases when the complexity of probe-probe intersection and limited numerical accuracy make the analytic method inapplicable. If that happens *a*_*p*_ is estimated using uniform grid-division of the surface of a unit sphere. The errors introduced by such an estimation are < 5% and thus should have no discernible impacts on our analysis of SES-defined electric and geometric properties.

Interior SES layers are computed as follows. After the exterior layer has been identified via the SES computation of a whole protein, it is removed for the computation of the first interior SSE layer, then the first interior SSE layer is removed for the computation of the second interior SSE layer, and the process stops when the number of atoms buried by the newly-computed SES layer is less than one hundred. In such a manner a whole protein is divided into a series of SES layers plus the innermost core.

### 2.4 SES-defined electric and geometric properties

Several electric and geometric properties have been defined in terms of atomic SES area and atomic partial chage to evaluate their relevance to protein-solvent interaction.

Let 𝔸 be a set of atoms in a protein. To accessible atom *i* ∈ 𝔸 we assign a nonzero SES area *a*(*i*):

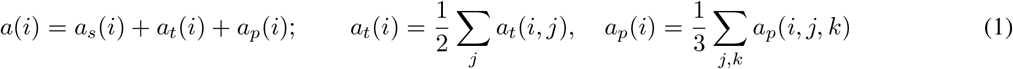

where *a*_*s*_(*i*), *a*_*t*_(*i*) and *a*_*p*_(*i*) are respectively the SAS, torus and probe areas for atom *i, a*_*t*_(*i, j*) is the area of a patch on a torus determined by atoms *i* and *j* ∈ *𝔸*, and *a*_*p*_(*i, j, k*) the area of a spherical polygon on a probe determined by atom *i* and atoms *j, k* ∈ *𝔸*. SAS area *a*_*s*_ is proportional to the square of atomic radius. Given the atomic SES areas set 𝔸 could be divided into a subset 𝕊 of surface atoms (or an SES layer of surface atoms) with nonzero areas and a subset 𝔹 of buried atoms with zero areas (or a core 𝔹 composed of the atoms with zero areas). From *a*_*s*_(*i*) and *a*_*t*_(*i*) + *a*_*p*_(*i*) we define a concave-convex ratio *r*_*cc*_(*i*) to estimate the local smoothness (or the extent of the exposure) of surface atom *i*. A surface atom with a low *r*_*cc*_ value means that a large portion of its atomic sphere is exposed and thus its local region is rugged. The average of the *r*_*cc*_(*i*)s for a set of atoms (e.g. 𝕊) is defined as *γ*_*cc*_.

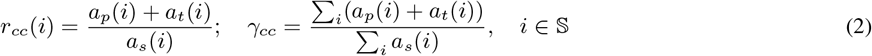

The average of the *γ*_*cc*_s over a set of structures is denoted as 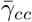.

On 𝕊 we define as follows its SES area *A*_*s*_ and area per atom *η*_*s*_, net surface charge *Q*_*s*_ and charge per atom *ρ*_*s*_.

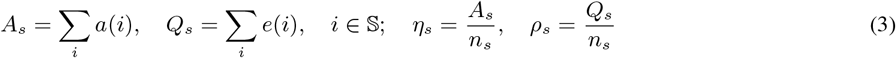

where *n*_*s*_ = |𝕊 | is the number of surface atoms in 𝔸 and *e*(*i*) the CHARMM partial charge [31] for atom *i*. In 𝔹 we define its net charge *Q*_*b*_ and charge per atom *ρ*_*b*_ as follows.

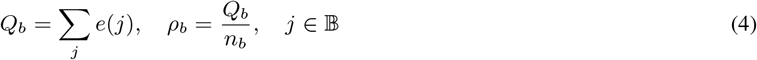

where *n*_*b*_ =| 𝔹 | is the number of buried atoms in 𝔸. The net charge *Q*, and charge per atom *ρ* for a *whole* protein are defined as follows.

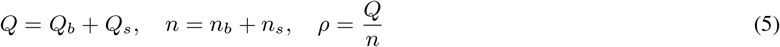

where *n* =| 𝔸 | is the total number of atoms in 𝔸. The averages of the *ρ*_*s*_s, *η*_*s*_s, *ρ*_*b*_s and *ρ*s over a set of structures will be denoted respectively as 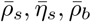 and 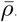.

The atoms in 𝕊 are further divided into four different subsets: a set of polar atoms **S**_*p*_, a set of positively-charged atoms **S**_+_, a set of negatively-charged atoms **S**_*−*_ and a set of apolar atoms **S**_*ap*_ (section S2 of SI), that is, 𝕊 = **S**_*ap*_ ∪ **S**_+_ ∪ **S**_*−*_ ∪ **S**_*p*_. Set **S**_+_ is composed of the positively-charged protons of the *ϵ*-ammonium group of a lysine residue and the guanidino group of an arginine residue. Set **S**_*−*_ is composed only of the side-chain oxygen atoms of both aspartic acid and glutaric acid residues. The atoms in **S**_+_, **S**_*−*_ and **S**_*p*_ are either an H-bond donor or an H-bond acceptor. Their union ℍ = **S**_+_ ∪ **S**_*−*_ ∪**S**_*p*_ is a set of atoms capable of forming an H-bond with other atoms. In a protein structure some of the atoms in ℍ may not form any detectable H-bond with other atoms, so ℍ is further divided into two subsets ℍ = **H**_b_ ∪ **H**_nb_ where the atoms in **H**_b_ form at least one H-bond with a protein atom while those in **H**_nb_ form none. A threshold of −555.55 for H-bond energy computed according to a formula in DSSP [32] is used to determine whether or not an atom in ℍ forms any H-bond with another protein atom. The SES areas per atom on these sets are defined as follows.

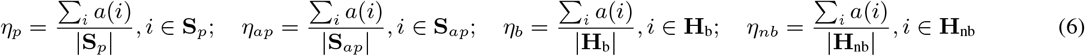

where two vertical bars denote the cardinality of a set. Their averages over a set of structures (e.g. 𝕊 ∈ 𝕄) are denoted respectively as 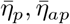 and 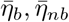. To characterize set ℍ we introduce a ratio *R*_h_ defined as 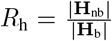. The average of *R*_h_s over a set of structures will be denoted as 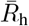.

One key feature of an extended conformation generated by CNS is that the side chain of each residue is generally much more exposed than the corresponding one in the native state. The extended conformations likely do not represent any real unfolded state. They are used here only as references for SES-defined electric and geometric properties.

## 3 Results and Discussion

In this section we begin with several electric and geometric properties defined in terms of atomic SES area, atomic partial charge and protein atom type for both exterior layer and interior layer, and then discuss their possible relevance to protein-solvent interaction. With the assignment to each atom the three different types of SES areas, partial charge and H-bonding capability a variety of quantities could be defined. In this paper we only present those that are likely related to best adaptation of a soluble protein to aqueous solvent.

### 3.1 The electric property of exterior SES layer and interior SES layer

Previous studies of protein surfaces, mainly SAS and VDW surface and occasionally SES, have shown that polar residues especially charged ones prefer to be on the surface of a soluble protein [22, 23, 24, 25]. In principle protein-solvent interaction is electrostatic in nature [14]. In theory surface charge and dipole moment are thought to be important for protein solvation [9, 10, 11, 13]. With these in mind in this section we first analyze atomic charge distribution in both exterior layer and interior layer for the soluble monomers in 𝕄. The analysis shows that not only the *ρ*_*s*_s of the exterior layers are all *negative* but the *ρ*_*s*_s for the successive SES layers of each structure change in a *zigzag* fashion from exterior to the interior and with a trend towards more positive. We then discuss the significance of such a multilayered organization to protein-solvent interaction in particular and protein structure in general.

#### 3.1.1 The net partial charges of the exterior layers and of the cores of water-soluble proteins

As shown in Fig. 1 all the structures in 𝕄 have a net negative surface charge (negative *Q*_*s*_ and *ρ*_*s*_). Moreover *ρ*_*s*_ decreases very slowly with increasing protein size and becomes essentially independent of protein size when number of surface atoms reaches about 2, 000. The near charge neutrality of a protein requires the net partial charges of the corresponding core to be positive. Indeed the *ρ*_*b*_ for every structure in 𝕄 is positive (Fig. 1) and *ρ*_*b*_ also decreases very slowly with protein size and becomes essentially independent of protein size when number of surface atoms reaches about 2, 000. Furthermore the 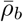 and the 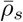 for 𝕄 have very close magnitudes and so do the *ρ*_*s*_ and the *ρ*_*b*_ for each individual structure. For reference the 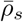 for 𝕌 is positive while the 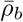 for 𝕌 is negative (Table S1 of SI). The buried backbone nitrogen and oxygen atoms account largely for the negativity of the 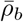 for 𝕌.

**Figure 1:**
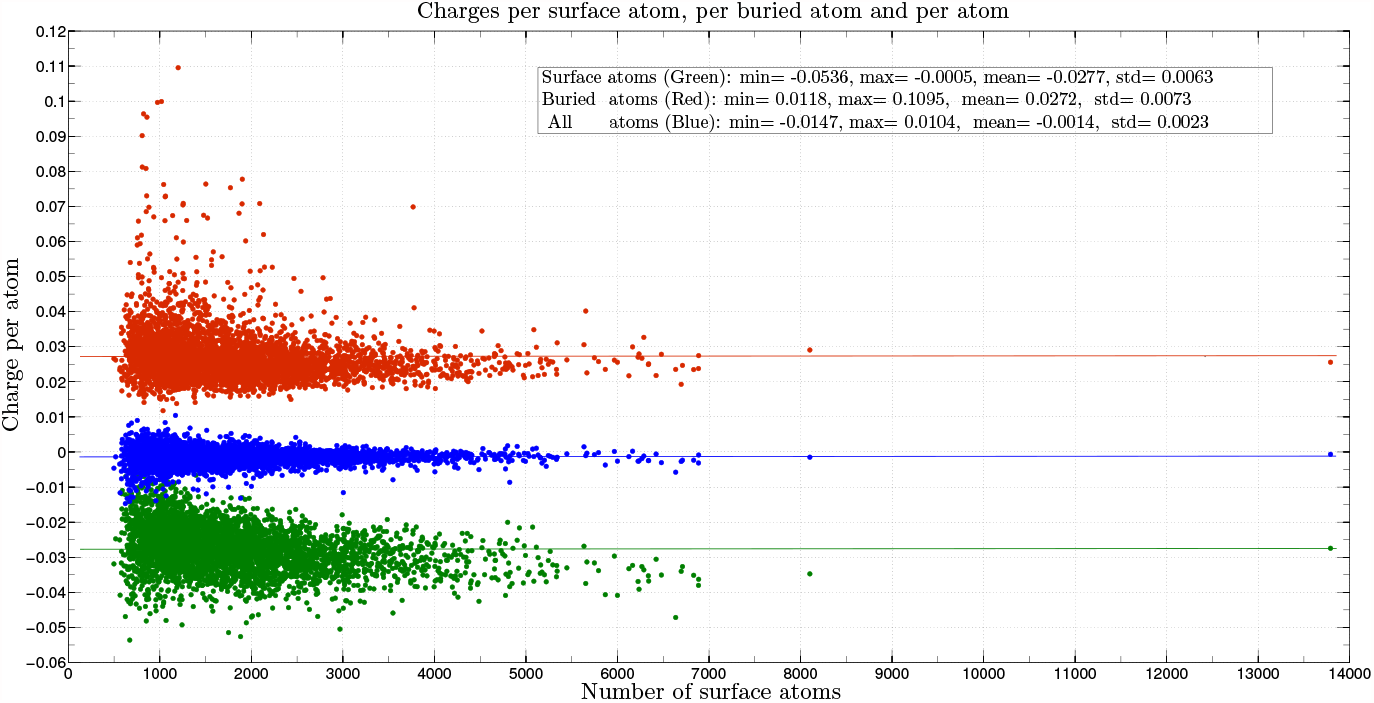
The *ρ*_*s*_s, *ρ*_*b*_s and *ρ*s for 𝕄. Their respective means as depicted by the three lines are 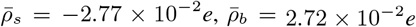, and 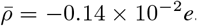. The deviations from zero of the *ρ*s may be explained as follows. In CHARMM force field the positively-charged residues, Arg, Lys and Hsp, have a net charge of +1*e* while the negatively-charged residues, Asp and Glu, have a net charge of *−* 1*e*. In addition a PDB file may lack coordinates for some atoms. The x-axis is the number of surface atoms in a structure. The y-axis is charge per atom with a unit of *e*.

With some exceptions [33, 34] previous studies of protein surface [22, 23, 24, 25] are largely performed at *residue* level and one of important findings is that polar residues especially charged ones prefer to be on surface while apolar residues prefer to be buried inside. In the contrary we characterize surface in terms of geometric and electric properties defined at *atomic* level. The latter is more natural from the viewpoint of physics while the former is more suitable for linking surface property with biological function. Our analysis shows that folding into a native state in aqueous solution turns a protein into some sort of dipole with a net positive charge buried inside and a net negative charge exposed on the surface to possibly maximize its electrostatic attraction to solvent molecules [7, 35]. In other words, a soluble protein behaves, on average and as far as surface charge is concerned, as a micelle [36] with an exterior formed predominately by atoms with negative partial charge and an interior composed mainly of atoms with positive partial charge.

#### 3.1.2 The zigzag and layered charge distribution from exterior to the interior

As described above like a dipole the exterior layer of a water-soluble monomer has a net negative charge while its interior has a net positive charge. However it is not clear how the atomic partial charges are distributed inside a protein. To study the distribution we compute SES for successive interior layers. As shown in Figs. 2a and 2b and Tables 1 and 2 for each individual monomer in 𝕄 the *ρ*_*s*_ for the first interior SES layer has almost the same magnitude as the *ρ*_*s*_ for the exterior layer but with a *positive* sign. The *ρ*_*s*_ of the second interior SES layer is more negative than that of the first interior layer but more positive than that of the exterior layer. The *ρ*_*s*_ of the third interior SES layer is in general more positive than that of the second interior layer. The *ρ*_*s*_s of the last (innermost) interior SES layers vary more than those of the previous layers and may be less positive than that for the second last interior layer. The large variance in the *ρ*_*s*_s for the last interior layers is due mainly to the large variance in the numbers of atoms in the last layers. In addition for each monomer in 𝕄, *ρ*_*s*_ changes the most from the exterior layer to the first interior layer. Furthermore from the exterior to the interior of any monomer in 𝕄 the overall trend in *ρ*_*s*_ value is towards more positive. The trend of becoming more positive is also clear when one compares the change in *ρ*_*b*_ from the exterior to the interior of a soluble monomer (Figure S1 of SI).

**Table 1:** The 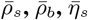 and 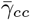 for 1,154 structures with four interior SES layers. The 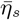 has a unit Å^2^, the 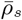 and 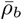 have a unit of *e*. The two values in each cell are respectively mean and standard deviation.

| Layer | $\bar{\rho}_s$ | $\bar{\rho}_b$ | $\bar{\eta}_s$ | $\bar{\gamma}_{cc}$ |
| --- | --- | --- | --- | --- |
| **Surface** | **-0.0306 (0.0053)** | **0.0234 (0.0030)** | **5.8304 (0.1104)** | **3.0569 (0.1932)** |
| 1st | 0.0313 (0.0073) | 0.0169 (0.0037) | 8.5995 (0.2625) | 2.6939 (0.2068) |
| 2nd | 0.0025 (0.0076) | 0.0374 (0.0072) | 9.1814 (0.3358) | 2.6674 (0.1727) |
| 3rd | 0.0324 (0.0096) | 0.0525 (0.0161) | 9.9651 (0.5783) | 2.2825 (0.1729) |
| 4th | 0.0406 (0.0181) | 0.1094 (0.0446) | 11.4822 (1.1141) | 1.7047 (0.2744) |

**Table 2:** The 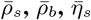 and 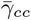 for 37 structures with five interior SES layers. The 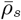s for the structures with five interior layers are more negative than the 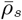s for those with four interior layers (Table 1). The number of layers increases with protein size while the 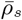 for a set of exterior layers becomes more negative with increasing number of layers. By comparison the 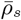 for all the structures in 𝕄 is −0.0277*e* (Fig. 1), less negative than that for the set of structures with either four or five interior layers. The values in each cell and their units are the same as those in Table 1.

| Layer | $\bar{\rho}_s$ | $\bar{\rho}_b$ | $\bar{\eta}_s$ | $\bar{\gamma}_{cc}$ |
| --- | --- | --- | --- | --- |
| **Surface** | **-0.0328 (0.0051)** | **0.0209 (0.0024)** | **5.8286 (0.0960)** | **3.1590 (0.2264)** |
| 1st | 0.0313 (0.0063) | 0.0138 (0.0022) | 8.4455 (0.2341) | 2.8166 (0.1988) |
| 2nd | -0.0011 (0.0068) | 0.0295 (0.0054) | 8.9017 (0.2725) | 2.8098 (0.1399) |
| 3rd | 0.0266 (0.0092) | 0.0344 (0.0085) | 9.4007 (0.5229) | 2.4948 (0.1489) |
| 4th | 0.0260 (0.0125) | 0.0546 (0.0156) | 9.9181 (0.6511) | 2.0468 (0.1524) |
| 5th | 0.0393 (0.0209) | 0.1070 (0.0459) | 11.2604 (1.2122) | 1.7312 (0.2166) |

**Figure 2:**
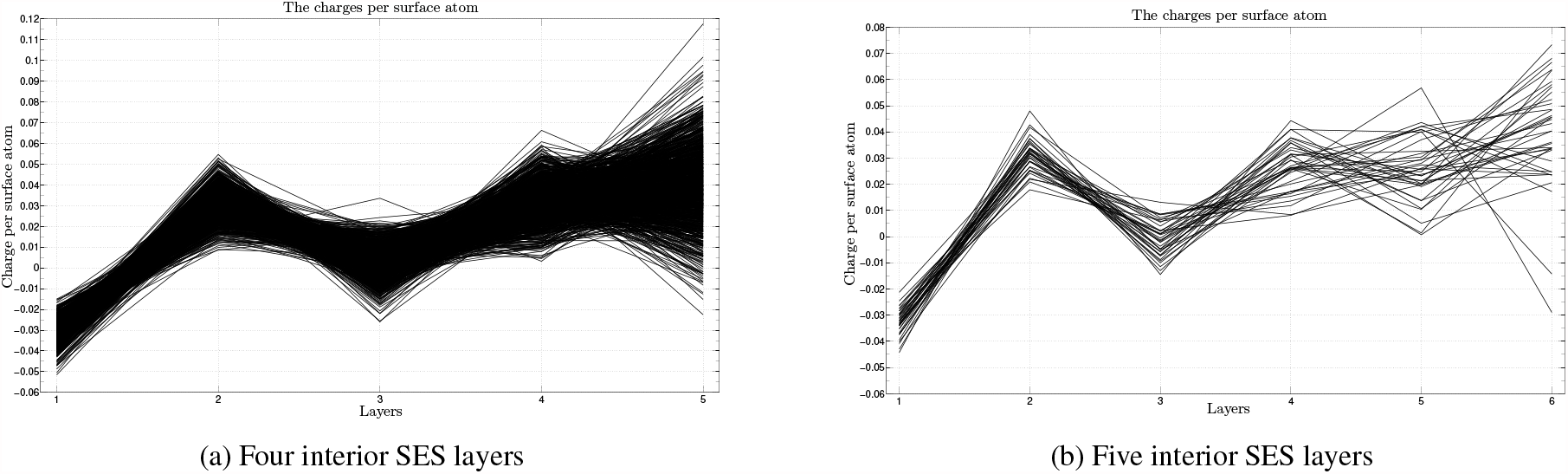
The *ρ*_*s*_s of exterior layer and interior SES layers. Figure (a) depicts the *ρ*_*s*_s of the 1, 154 structures with four interior SES layers while (b) depicts the *ρ*_*s*_s of the 37 structures with five interior layers. Other monomers in 𝕄 have only two or three interior SES layers and none of the monomers has more than five interior layers. The x-axis is layer index. The y-axis is charge per atom with a unit of *e*.

#### 3.1.3 The significance to protein-solvent interaction of layered and zigzag charge distribution

To the best of our knowledge we have not known any report of such a zigzag increase of net charge from the exterior to the interior of any protein except for a model of alternative layers of negative and positive charges alluded previously in MD simulation [37]. Both the steep increase from exterior SES layer to first interior SES layer and the zigzag pattern are expected to be closely related to protein-solvent interaction. Due to the importance of electric interaction to protein solvation, folding and structure, the steep increase and the zigzag pattern may give clue to the best adaptation of soluble proteins to aqueous solvent. In addition the non-uniform charge distribution from exterior to the interior suggests that there likely exists a similar one for dielectric constant [38].

### 3.2 The geometric properties of exterior SES layer and interior SES layer

The atomic SES areas of a protein could be used not only to separate its atoms into a set of surface atoms and a set of buried atoms but the three composing areas, *a*_*s*_, *a*_*t*_ and *a*_*p*_, of a surface atom could also be used to define atomic geometric properties such as *r*_*cc*_ and *γ*_*cc*_ to describe its local geometry. In the following we first present the SES areas per atom (*η*_*s*_s) and the concave-convex ratios (*γ*_*cc*_s) for both the exterior layers and the interior layers of the soluble monomers in 𝕄. We then show that inside each monomer from exterior to the interior there exists a multilayered organization in terms of both *η*_*s*_ and *γ*_*cc*_. Such a multilayered organization is expected to be relevant to protein-solvent interaction.

#### 3.2.1 The *η*_*s*_ and *γ*_*cc*_ of exterior SES layer

The *η*_*s*_s for all the structures in 𝕄 have an average 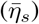 of 5.861Å^2^ and a narrow range with a standard deviation of 0.133Å^2^ (Table 3). Moreover except for rare cases *η*_*s*_ increases linearly with the number of surface atoms in a structure. The *γ*_*cc*_s for all the structures in 𝕄 have an average 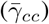 of 2.851 and a standard deviation of 0.247. For reference the 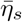 for 𝕌 is 1.1-fold larger than that for 𝔽 while the 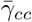 for 𝕌 is 1.6-fold smaller than that for 𝔽 (Table S2 of SI). The differences in *η*_*s*_ and *γ*_*cc*_ between the folded and the extended conformations suggest that folding into a native state smooths out the roughness of the local region surrounding an exposed atom.

**Table 3:** The average *η*s and *γ*_*cc*_s for all surface atoms (set 𝕊), apolar atoms (set S_*ap*_) and polar atoms (set S_*p*_), positively-charged atoms (set S_+_), negatively-charged atoms (set S_*−*_) and H-bond donors and acceptors that form either H-bond (set H_b_) or no H-bond with protein atoms (set H_nb_). The two numbers in each cell are respectively mean and standard deviation. They are computed over the exterior layers for all the monomers in 𝕄. The 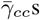 in Tables 1 and 2 are for two subsets in 𝕄 that have respectively four and five interior SES layers. The 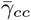 for all surface atoms in this table is for all the structures in 𝕄 that also include structures with two and three interior SES layers.

|  | S | S<sub>ap</sub> | S<sub>p</sub> | S<sub>+</sub> | S<sub>-</sub> | H<sub>b</sub> | H<sub>nb</sub> |
| --- | --- | --- | --- | --- | --- | --- | --- |
| $\eta$ | 5.861 (0.133) | 4.812 (0.127) | 7.750 (0.331) | 6.080 (0.323) | 15.023 (1.037) | 6.897 (0.460) | 9.085 (0.439) |
| $\gamma_{cc}$ | 2.851 (0.241) | 3.490 (0.347) | 2.686 (0.296) | 1.891 (0.358) | 1.370 (0.255) | 3.151 (0.496) | 1.983 (0.226) |

#### 3.2.2 The *η*_*s*_ and *γ*_*cc*_ of interior SES layer

The *η*_*s*_ for a soluble monomer in 𝕄 increases invariably from exterior to the interior (Figs. 3a, 3b). Furthermore from exterior SES layer to first interior SES layer 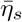 increases 1.4-fold, and this rate of increase is much larger than that between any two successive interior SES layers. In the contrary the *γ*_*cc*_s for the monomers decrease from exterior to the interior (Figs. 4a, 4b). Specifically except for a small number of monomers in 𝕄 that the *γ*_*cc*_s between two consecutive interior SES layers change in a zigzag manner, the overall trend in *γ*_*cc*_ is towards reduction. Taken together the increase in *η*_*s*_ and the decrease in *γ*_*cc*_ show that the interior layer of a soluble monomer is locally more rugged than the exterior layer. An interior SES layer is created by removing the previous layer of accessible protein atoms while the atoms in an exterior layer interact directly with solvent molecules. The atoms in the former are optimized to interact with other protein atoms while the local geometry of the latter must have at least been partially optimized for protein-solvent interaction.

**Figure 3:**
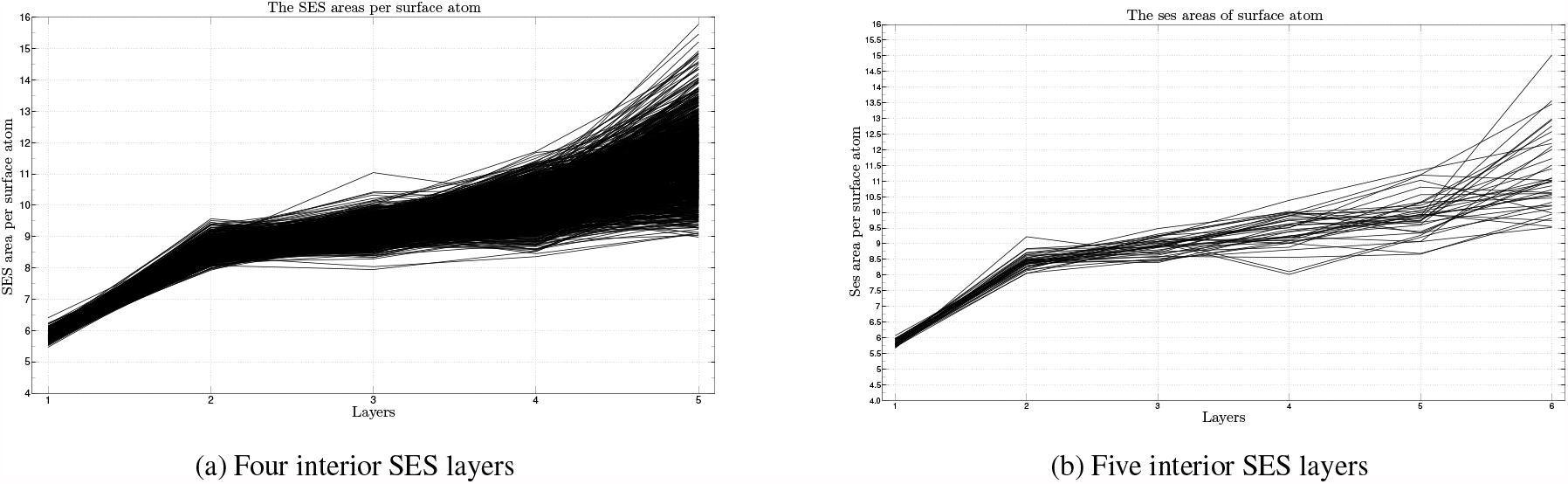
The *η*_*s*_s of exterior SES layer and interior SES layer. Figure (a) depicts the *η*_*s*_s for the 1, 154 structures with four interior layers while (b) depicts the *η*_*s*_s for the 37 structures with five interior layers. The x-axis is layer index. The y-axis is SES area per atom with a unit of Å^2^.

**Figure 4:**
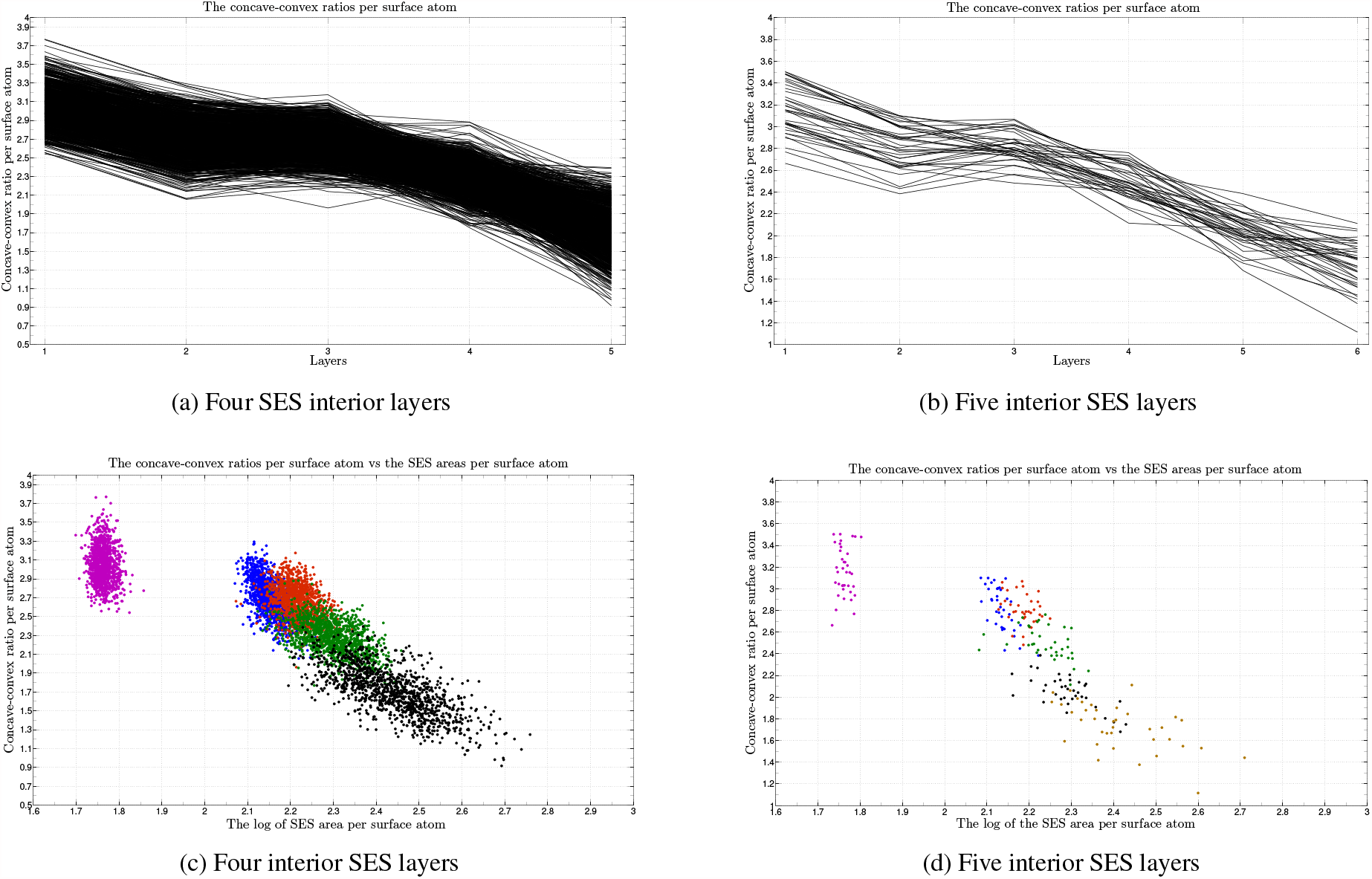
The *γ*_*cc*_s of exterior SES layer and interior SES layer. Figure (a) depicts the *γ*_*cc*_s for the 1, 154 structures with four interior SES layers while (b) depicts the *γ*_*cc*_s for the 37 structures with five layers. In figure (a, b) the x-axis is layer index while the y-axis is *γ*_*cc*_. In figure (c) the *γ*_*cc*_s for the exterior layers and the first to the fourth interior layers are colored respectively in magenta, blue, red, dark green and black. In figure (d) the *γ*_*cc*_s for the exterior layer and the first to the fifth interior layers are colored respectively in magenta, blue, red, dark green, black and orange. The larger variance in *γ*_*cc*_ for more inner layers is due mainly to the large difference in the number of atoms in each layer. The x-axis is in figures (c) and (d) is log *η*_*s*_ where *η*_*s*_ is SES area per atom.

The difference between exterior and interior SES layers is more striking if we plot *γ*_*cc*_ verse log *η*_*s*_, the logarithm of SES area per atom. As shown in Figs. 4c and 4d, all the *γ*_*cc*_s for the exterior layers are well separated from those for the interior layers. Furthermore unlike the *γ*_*cc*_ for an interior layer the *γ*_*cc*_ for an exterior layer does not change with either number of surface atoms or SES area. In the contrary the *γ*_*cc*_s for two neighboring interior SES layers overlap with each other and *γ*_*cc*_ decreases continuously from the first interior layer to the last interior layer. The well-separation of the exterior layers from the interior layers in terms of *γ*_*cc*_ and the continuous decrease in *γ*_*cc*_ from exterior to the interior show that *γ*_*cc*_ is a geometric property relevant to protein-solvent interaction. Furthermore the continuous increase in *η*_*s*_ together with the continuous decrease in *γ*_*cc*_ from exterior to the interior are consistent with the previous observation that the core of a soluble protein is tightly packed [26]. However previous structural analyses focused mainly on the difference in packing between protein surface [39] and protein core [26] while the packing difference in different parts of a protein core has not been well documented. With the separation of a protein core into successive interior SES layers we are able to show that in terms of *η*_*s*_ and *γ*_*cc*_ not only the first core^9^ of a protein differ from the solvent-accessible surface but a more inner core also differs from a more outer core. Specifically from exterior to the interior the packing becomes increasingly tighter.

### 3.3 The multilayered organization of soluble monomers and its implications for protein solvation and structure

As described above for each monomer in 𝕄 in terms of *ρ*_*s*_, *ρ*_*b*_, *η*_*s*_ and *γ*_*cc*_ there exist not only differences between the exterior layer and the interior SES layers but also between two neighboring interior layers. Specifically from the exterior SES layer to the innermost interior SES layer of a monomer (1) *ρ*_*s*_ changes in a zigzag manner with its value most negative for the exterior layer and with a trend towards more positive, (2) *η*_*s*_ and *ρ*_*b*_ increase continuously but, (3) *γ*_*cc*_ decreases almost continuously. Thus in terms of these SES-defined electric and geometric properties the atoms in a soluble monomer are not distributed randomly but with a multilayered organization. The increasingly tight-packing from exterior to the interior as implied by the increase in *η*_*s*_ and the decrease in *γ*_*cc*_ together with the increase in net positive charge from exterior to the interior mean that electric repulsion becomes stronger with more interior layers. Thus the multilayered organization may set an upper limit to the size and the overall shape of a soluble monomer. To satisfy such a multilayered organization a soluble monomer either adopts a non-spherical shape or limits the number of interior layers for a spherically-shaped protein. In fact out of the 4, 290 soluble monomers with up to 27, 859 atoms only 37 monomers have five interior SES layers and none has six or more interior layers. Such a multilayered organization has likely been evolved to achieve an optimal interaction with aqueous solvent. This layered organization of a soluble protein is an extension of its hydration layers [40]. The combined multilayered organization consisting of several hydration layers, the exterior layer and several interior layers should be a general feature of a protein-solvent system and thus should play a vital role in the solvation, folding and structure of water-soluble proteins.

A multilayered organization for proteins has been described previously using atom depth [41]. However there exist important differences between the layered organization defined by SES and that defined by atom depth. Firstly the layers computed analytically by our SES algorithm differ from the layers computed by atom depth. Secondly atom-depth based analysis focused only on the number of atoms in a layer while our analysis focuses on layer’s SES-defined electric and geometric properties. Finally atom-depth based analysis has revealed no relationship between layered organization and protein-solvent interaction while our analysis shows that the SES-derived multilayered organization is a general feature of a protein-solvent system.

### 3.4 The SES areas per atom and the concave-convex ratios of apolar atoms and polar atoms and of the atoms forming H-bond and the atoms forming no H-bond

Previous structural analyses [22, 23, 24, 25] have found that at *residue* level polar residues especially charged ones prefer to be on the surface of a soluble protein while apolar residues prefer to be buried inside. The surface atoms in the exterior layer of a soluble protein interact directly with solvent molecules through intermolecular H-bonding [28] that is directional and intermolecular VDW attraction [27] that is nondirectional. Thus at atomic level the H-bond donors and acceptors in an exterior layer are expected to have geometric properties different from the other surface atoms. With atomic SES area and the separation of surface atoms into polar subset (**S**_*p*_) and apolar subset (**S**_*ap*_) it is possible to evaluate the preferences at *atomic* level using SES-defined geometric properties such as *η*^10^ and *γ*_*cc*_. Specifically in this section we analyze the preferences in the exterior layers of the soluble monomers in 𝕄 by comparing the *η*_*s*_s and *γ*_*cc*_s for apolar atoms and polar atoms and for H-bond donors and acceptors that form an H-bond with a protein atom and those that form no such an H-bond.

#### 3.4.1 The *η*s and *γ*_*cc*_s for the polar atoms and the apolar atoms and the charged atoms

The 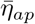 for the **S**_*ap*_s in 𝕄 is only 4.812Å^2^ while the 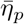 for the **S**_*p*_s in 𝕄 is 7.750Å^2^ (Fig. 5a and Table 3), the latter is 1.61-fold larger than the former. In other words on average the SES area of a polar atom is 1.61-fold larger than that of an apolar atom. For reference the same ratio for the 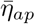 and 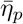 in 𝕌 is only 1.22 (Table S2 of SI). Set **S**_*p*_ does not include the negatively-charged side-chain oxygen atoms in Asp, Glu and the positively-charged protons in Lys and Arg. The 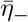 for the **S**_*−*_s in 𝕄 is 15.023Å^2^ and the 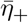 for the **S**_+_s in 𝕄 is 6.080Å^2^, both are much larger than 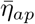^11^. Taken together it means that the H-bond donors and acceptors in an exterior layer on average have *η*_*s*_s larger than those for the accessible apolar atoms. For reference the 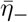 (17.738Å^2^) and the 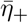 (6.927Å^2^) for 𝕌 are only slightly larger than those for 𝔽 (Table S2 of SI). In an extended conformation the side chain carboxyl groups of Asp and Glu, and the *ϵ*-ammonium group of Lys and the guanidino group of Arg are almost fully exposed. Thus in terms of SES area per atom the charged residues on the surface of a soluble monomer are close to be fully exposed. This confirms the preference of these charged residues for surface [22, 23, 24, 25]. The negatively-charged side-chain oxygen atoms are H-bond acceptor while the positively-charged protons are H-bond donor. As detailed later the large atomic SES areas for a donor and an acceptor are advantageous for them to form H-bonds with solvent molecules.

**Figure 5:**
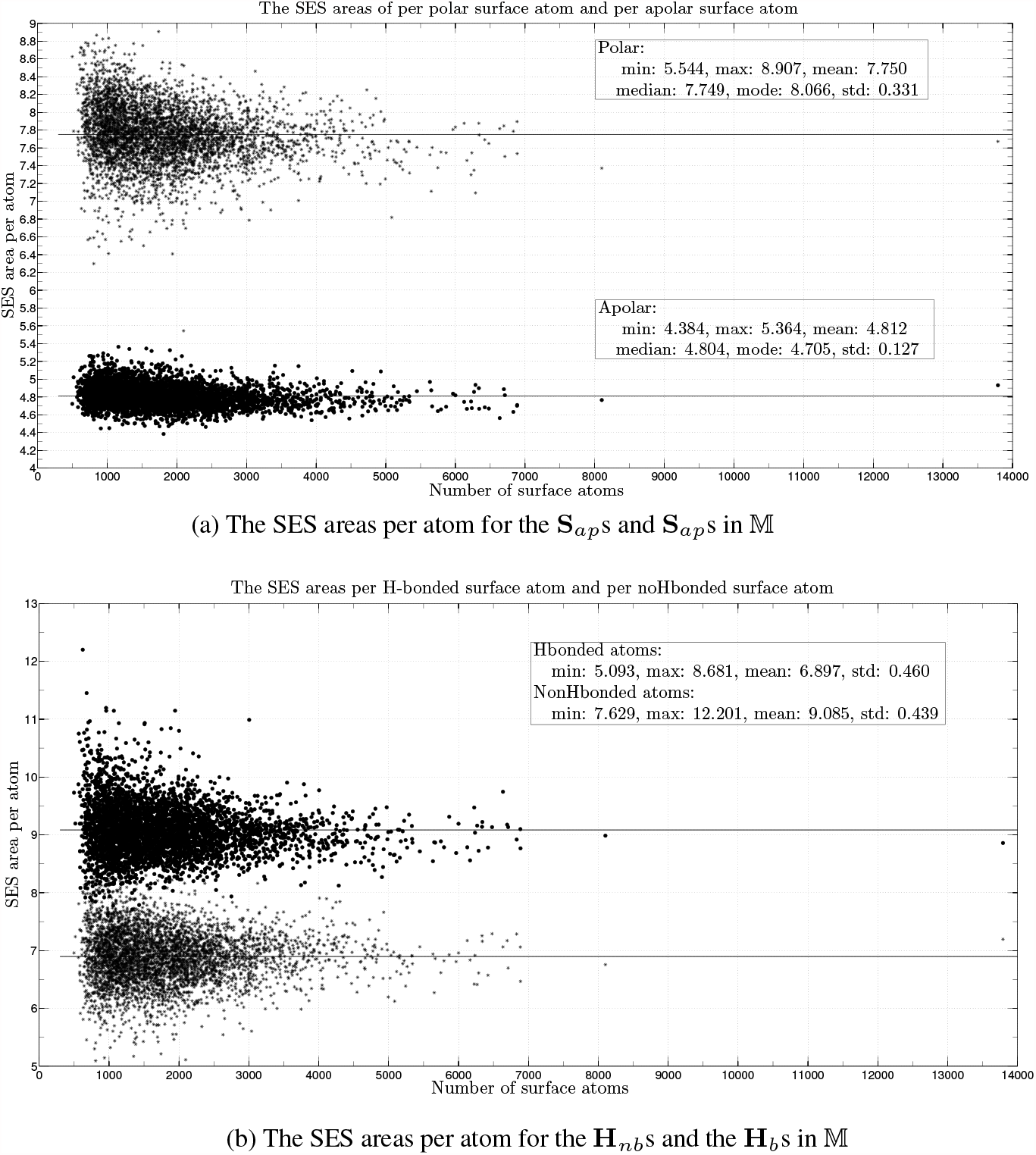
The *η*s for the S_*p*_s, S_*ap*_s and H_b_s, H_nb_s in 𝕄. The middle insert in each figure lists mean, median, mode and standard deviation with the mean depicted by a line. Figure (a) depicts the *η*s for the **S**_*p*_s and **S**_*ap*_s while figure (b) depicts the *η*s for the **H**_b_s and **H**_nb_s. The x-axis is the number of surface atoms in a structure and the y-axis is SES area per atom in Å^2^.

The 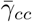 for the **S**_*ap*_s in 𝕄 is 3.490 and is 1.3-fold larger than the 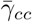 for the **S**_*p*_s (Table 3). The 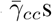 for the **S**_+_s and the **S**_*−*_s are even smaller than that for the **S**_*p*_s. For reference the 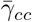 for the **S**_*ap*_s in 𝕌 is only 1.973 (Table S2 of SI). It is only slightly larger than that for the **S**_*p*_s in 𝕌, the latter is 1.862. The difference in 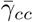 between the apolar atoms and the polar atoms in the exterior SES layer of a soluble monomer shows that the local region of an accessible apolar atom is on average smoother than that of an accessible polar atom. The apolar atoms in an exterior layer interact with solvent molecules through VDW attraction. A smooth local region for an accessible apolar atom likely optimizes its intermolecular VDW attraction with solvent molecules with a minimum disruption to water’s internal configuration. This is consistent with the importance of VDW attraction for the solvation of hydrophobic molecules in aqueous solvent [27].

#### 3.4.2 The *η*s and *γ*_*cc*_s for the H-bond donors and acceptors that form an H-bond and those that form no H-bond

Among a set of H-bond donors and acceptors (set ℍ) in either the exterior layer or the core of a soluble protein some form no H-bond with any protein atom. Thus the atoms in ℍ could be further divided into two subsets, that is, ℍ = **H**_b_ ∪ **H**_nb_ where the atoms in **H**_b_ form at least one H-bond with a protein atom while those in **H**_nb_ form none. The ratio 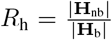 is a property for set ℍ. The 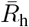 for the sets of surface atoms (set 𝕊s) in 𝕄 is 1.66 while the 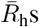 for the sets of buried atoms (set 𝔹s) is only 0.154. It means that on average about two-thirds of the H-bond donors and acceptors in the exterior layer of a soluble monomer form no H-bond with any protein atom. Most of them likely form H-bonds with solvent molecules. In the contrary less than one-sixths of the H-bond donors and acceptors in the corresponding core form no H-bond with any protein atom. For reference the 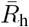 for the the 𝕊s in 𝕌 is 2.21 while the 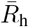 for the cores in 𝕌 is 1.64, these two values are much closer to each other than the corresponding pair of values for 𝕄^12^.

Furthermore as shown in Fig. 5b and Table 3 the 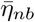 (9.085Å^2^) for the **H**_nb_s in 𝕄 is 1.32-fold larger than that for the **H**_b_s. However the 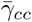 (1.983) for the **H**_nb_s in 𝕄 is 1.61-fold smaller than the 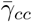 (3.153) for the **H**_b_s. For reference the 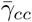 for the **H**_nb_s in 𝕌 is 1.424, only 1.26-fold smaller than the 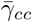 (1.789) for the **H**_b_s in 𝕌 (Table S2 of SI). Since the inter-atomic distance between two H-bonded atoms is smaller than the summation of their respective VDW radii, the larger SES area a donor or an acceptor has, the less likely a solvent molecule clashes with its neighboring protein atoms and less likely to perturb water’s H-bonded network when it forms an optimal H-bond with a water molecule. The large 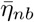 value and the small 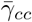 value together with the large 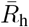 value for the **H**_nb_s in 𝕄 support the importance for protein solvation of the intermolecular H-bonding between accessible polar or charged atoms and solvent molecules [28].

#### 3.4.3 Both hydrogen bonding interaction and VDW attraction contribute to protein solvation

The differences in *η* and *γ*_*cc*_ between **S**_*ap*_ and **S**_*p*_ and between **H**_nb_ and **H**_b_ for soluble monomers show that these two SES-defined geometric properties are likely pertinent to protein solvation and structure via the optimization of both the intermolecular VDW attraction between accessible apolar atoms and solvent molecules and the intermolecular H-bonding interaction between polar and charged protein surface atoms and solvent molecules [42]. From an evolutionary perspective it seems that the surface of a naturally-occurring soluble monomer has been optimized for best interaction with aqueous solvent through optimal H-bonding interaction and optimal VDW attraction.

## 4 The possible limitations of the analysis

Our analysis relies on protein structures in crystalline state but soluble proteins function in solution state and their surfaces especially their active sites are flexible [43]. Furthermore, the surface of a crystal structure may be distorted by crystal packing though its interior is expected to be close to what exists in solution. It will be interesting to compare the SESs of NMR solution structures with the SESs of crystal structures. The analysis of SES-defined electric properties depends on the accuracy of atomic partial charge assigned by CHARMM. The values of the CHARMM atomic partial charges for a residue do not take into account whether or not the residue is on the surface of a protein or buried inside. It is possible that the polarization of the same atom in the same type of amino acid may depend on its micro-environment [44] and thus different values should be used depending on where it is in the protein. From this viewpoint the SES-defined geometric properties particularly those of interior layers are more reliable. SES itself is more a mathematical model than a truly physical model. It is generated by modeling both the protein atoms and water molecules as a sphere. Thus the conclusions reached by analyzing the SES-defined geometric and electric properties for soluble monomers are qualitative in nature.

## Supporting Information

### S1 The sequence similarity of the set of soluble monomers

There are 4, 290 structures in 𝕄. When downloaded from the PDB we selected a criterion that requires their sequence identities be less than 70%. To further evaluate their sequence similarity we run BLASTp [1] on all the pairs of structures that have < 15.0% differences in number of residues. Out of all such pairs there are 2, 983 pairs that have a BLASTp *Expect value* ≤1*e* − 10. Out of the 2, 983 pairs there are only 68 pairs that have a BLASTp *Positives value* ≥ 80.0%. The statistics for BLASTp Positives value for the 68 pairs are the following: min= 80.1%, max= 100.0%, mean= 86.4%, std = 6.1%. The three pairs, 1G8F–1R6X, 1H6T–2Y5Q and 1PRZ–1V9F, that have a BLASTp Positives value of 100.0% have respectively 511–395, 291–362 and 252–325 residues. The statistics for BLASTp Positives value for all the 2, 983 pairs are min= 35.4%, max= 100.0%, mean= 54.1%, std = 11.1%.

### S2 The apolar atoms and polar atoms and the positively-charged atoms and the negatively-charged atoms

The atoms in a protein are divided into four mutually-exclusive types: *apolar, polar, negatively-charged and positively-charged*.

The type of polar atoms is: {HN, N, O, HT1, HT2, HT3, OT2, OT1, K_NZ, R_NE, R_NH1, R_NH2, N_OD1, N_ND2, N_HD21, N_HD22, Q_OE1, Q_NE2, Q_HE21, Q_HE22, HSE_ND1, HSE_HD1, HSE_NE2, HSE_HE2, HSD_ND1, HSD_HD1, HSD_NE2, HSD_HE2, HSP_ND1, HSP_HD1, HSP_NE2, HSP_HE2, H_ND1, H_HD1, H_NE2, H_HE2, S_OG, S_HG1, T_OG1, T_HG1, Y_OH, Y_HH, W_NE1, W_HE1 }.

The type of negatively-charged atoms is: {D_OD1, D_OD2, E_OE1, E_OE2}.

The type of positively-charged atoms is: {K_HZ1, K_HZ2, K_HZ3, R_HE, R_HH11, R_HH12, R_HH21, R_HH22 }. The type of apolar atoms includes all the atoms other than the above three types.

The name of each atom consists of two parts separated by a hyphen: the part before the hyphen is residue name while the part after the hyphen is atom name in CHARMM nomenclature. Each polar or charged atom is either a hydrogen bond (H-bond) donor or an H-bond acceptor.

### S3 The charges per atom and SES areas per atom for both the extended conformations and folded conformations

Table S1 lists the means, 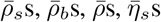 and 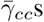, and the standard deviations for the *ρ*_*s*_s, *ρ*_*b*_s, *ρ*s, *η*_*s*_s and *γ*_*cc*_s of both the extended conformations in 𝕌 and their corresponding folded conformations in 𝔽.

**Table S1:** The *ρ*_*s*_s, *ρ*_*b*_s, *ρ*s, *η*_*s*_s and *γ*_*cc*_s of both the extended conformations (𝕌) and folded conformations (𝔽). The two values in each cell are respectively mean and standard deviation.

| | $\rho_s (\times 10^{-3}e)$ | $\rho_b (\times 10^{-3}e)$ | $\rho (\times 10^{-3}e)$ | $\eta_s (\text{\AA}^2)$ | $\gamma_{cc}$ |
| --- | --- | --- | --- | --- | --- |
| $\mathbb{U}$ | +0.516 ( 2.91 ) | -7.31 ( 32.33 ) | 0.235 ( 2.54 ) | 6.466 ( 0.101 ) | 1.786 ( 0.027 ) |
| $\mathbb{F}$ | -26.56 ( 6.23 ) | +27.58 ( 7.74 ) | -0.125 ( 2.42 ) | 5.862 ( 0.132 ) | 2.809 ( 0.233 ) |

### S4 The layered distribution of charge per buried atom for soluble monomers

Figure S1 shows the layered distribution of *η*_*b*_s for the 1, 154 structures with four interior SES layers and the 37 structures with five interior layers. Except for the slight reduction from the exterior layers to the first interior layers for all the other successive interior layers the trend is towards more positive.

**Figure S1:**
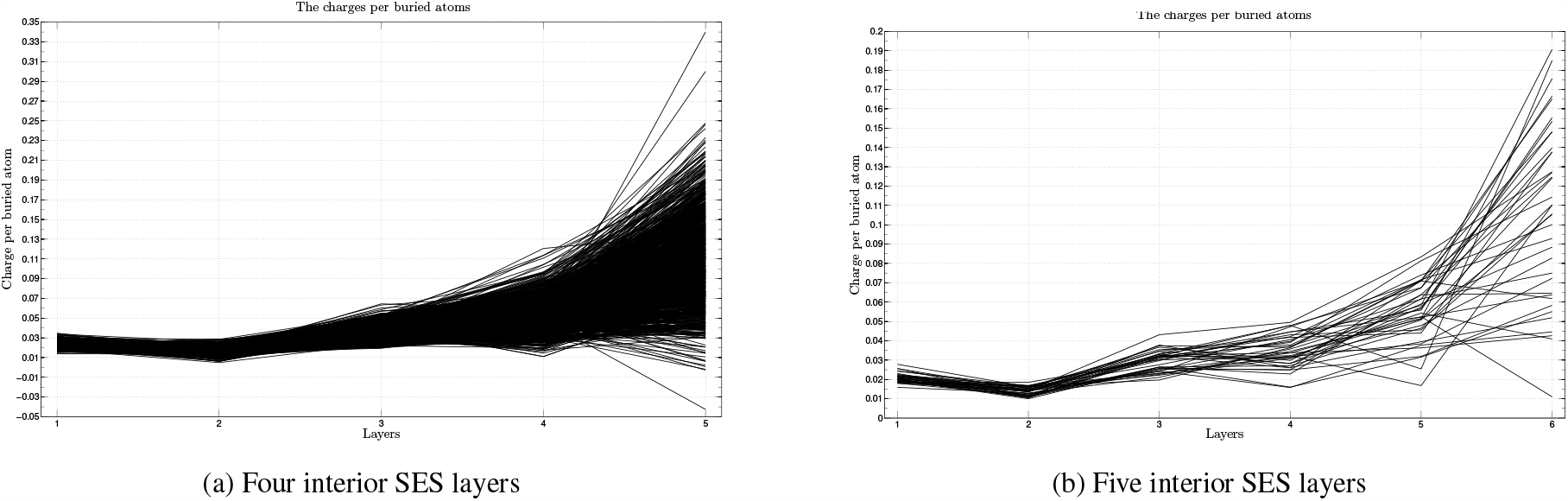
The *ρ*_*b*_s of exterior layer and interior SES layers. Figure (a) depicts the *ρ*_*b*_s of the 1, 154 structures with four interior SES layers while (b) depicts the *ρ*_*b*_s of the 37 structures with five interior layers. Other monomers in 𝕄 have only two or three interior SES layers. The x-axis is layer index. The y-axis is charge per atom with a unit of *e*.

### S5 The SES areas per atom and the concave-convex ratios per atom for both the extended conformations and folded conformations

Table S2 lists the means, 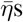 and 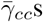, and the standard deviations for the *η*s and *γ*_*cc*_s of both the extended conformations in 𝔽 and their corresponding folded conformations in 𝕌. The means and standard deviations are computed for seven different sets (𝕊, **S**_*ap*_**S**_*p*_, **S**_+_, **S**_*−*_, **H**_b_ and **H**_nb_) of surface atoms in the exterior layers of both 𝕌 and 𝔽.

**Table S2:** The average *η*s and *γ*_*cc*_s for all surface atoms (set 𝕊), apolar atoms (set S_*ap*_) and polar atoms (set S_*p*_), positively-charged atoms (set S_+_) and negatively-charged atoms (set S_*−*_), and H-bond donors and acceptors that form either H-bond (set H_b_) or no H-bond with protein atoms (set H_nb_). The two numbers in each cell are mean and standard deviation. They are computed over the exterior layers of all the extended conformations in 𝕌 and the exterior layers of all the folded conformations in 𝔽.

| | $\mathbb{S}$ | $\mathbb{S}_{ap}$ | $\mathbb{S}_p$ | $\mathbb{S}_+$ | $\mathbb{S}_-$ | $\mathbb{H}_b$ | $\mathbb{H}_{nb}$ |
| --- | --- | --- | --- | --- | --- | --- | --- |
| $\mathbb{U} : \eta$ | 6.466 (0.101) | 5.585 (0.089) | 8.150 (0.204) | 6.927 | 17.738 | 8.361 (0.388) | 8.862 (0.247) |
| $\mathbb{F} : \eta$ | 5.862 (0.132) | 4.807 (0.119) | 7.758 (0.347) | 6.068 | 15.172 | 6.914 (0.476) | 9.080 (0.454) |
| $\mathbb{U} : \gamma_{cc}$ | 1.786 (0.027) | 1.973 (0.045) | 1.862 (0.090) | 1.830 | 1.339 | 1.789 (0.191) | 1.424 (0.086) |
| $\mathbb{F} : \gamma_{cc}$ | 2.809 (0.233) | 3.475 (0.333) | 2.629 (0.275) | 0.894 | 0.682 | 3.083 (0.469) | 1.933 (0.219) |

## ^1^Abbreviations

SES: solvent-excluded surface
SAS: solvent accessible surface
VDW: van der Waals
H-bond: hydrogen bond
H-bonding: hydrogen bonding
2D: two-dimensional
MD: molecular simulation
PDB: Protein Data Bank
CHARMM: Chemistry at Harvard Macromolecular Mechanics
SI: Supporting Information

## Footnotes

2 In the rest of the paper, *solvent-accessible atom, accessible atom, surface atom and exposed atom* are used interchangeably.

3 No interior SES layers are computed for𝕌.

4 The SAS area, torus area and probe area for a layer of surface atoms are defined respectively as the summations of atomic SAS area, torus area and probe area over all the atoms in the layer.

5 For brevity the expression *“from exterior to the interior”* is used to mean either from the exterior SES layer to the innermost interior SES layer or from the first core of buried atoms to the last core of buried atoms of a soluble protein.

6 Each polar atom is either an H-bond donor or acceptor.

7 The difference in the number of atoms observed and the number of atoms expected from a protein sequence.

8 A gap in a protein chain means that either one or several consecutive interior residues have no ATOM statement in the PDB file.

9 The first core of a protein means the set of atoms buried by its exterior SES layer.

10 SES area per atom for a nonspecific set of surface atoms will be denoted as *η* without subscript.

11 A value of 6.080Å^2^ for 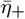 is in fact relatively large considering that the atomic radius of a proton is only 1.09Å and SAS area is proportional to the square of atomic radius.

12 The 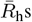 for 𝕌 are larger than the corresponding 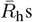 for 𝕄 since in an extended conformation the side chains are fully-exposed and there exists no H-bond between backbone amide protons and backbone carbonyl oxygens.

## References

[1] C. Tanford. Contribution of hydrophobic interactions to the stability of the globular conformation of proteins. Journal of the American Chemical Society, 84(20):4240–4247, 1962.

[2] W. Kauzmann. Thermodynamics of unfolding. Nature, 325:763–764, February 1987.

[3] T. Y. Lin and S. N. Timasheff. On the role of surface tension in the stabilization of globular proteins. Protein science, 5(2):372–381, 1996.

[4] Y. Levy and J. N. Onuchic. Water mediation in protein folding and molecular recognition. Annu. Rev. Biophys. Biomol. Struct., 35:389–415, 2006.

[5] M. Bellissent-Funel, A. Hassanali, M. Havenith, R. Henchman, P. Pohl, F. Sterpone, D. van der Spoel, Y. Xu, and A. E Garcia. Water determines the structure and dynamics of proteins. Chemical Reviews, 116(13):7673–7697, 2016. PMID: 27186992.

[6] W. Blokzijl and J. B. F. N. Engberts. Hydrophobic effects. opinions and facts. Angewandte Chemie International Edition in English, 32(11):1545–1579, 1993.

[7] R. M. Kramer, V. R. Shende, N Motl, C. N. Pace, and J. M. Scholtz. Toward a molecular understanding of protein solubility: Increased negative surface charge correlates with increased solubility. Biophysical Journal, 102(8):1907–1915, 2012.

[8] C. Højgaard, C. Kofoed, R. Espersen, K. E. Johansson, M. Villa, M. Willemoës, K. Lindorff-Larsen, K. Teilum, and J. R. Winther. A soluble, folded protein without charged amino acid residues. Biochemistry, 55(28):3949–3956, 2016. PMID: 27307139.

[9] L. Onsager. Electric moments of molecules in liquids. Journal of the American Chemical Society, 58(8):1486–1493, 1936.

[10] C. Tanford and J. G. Kirkwood. Theory of protein titration curves. i. general equations for impenetrable spheres. Journal of the American Chemical Society, 79(20):5333–5339, 1957.

[11] G. King and A. Warshel. A surface constrained all atom solvent model for effective simulations of polar solutions. The Journal of Chemical Physics, 91(6):3647– 3661, 1989.

[12] J. Chen, C. L. Brooks III, and J. Khandogin. Recent advances in implicit solvent-based methods for biomolecular simulations. Current Opinion in Structural Biology, 18(2):140 – 148, 2008. Theory and simulation / Macromolecular assemblages.

[13] A. V. Marenich, C. J. Cramer, and D. G. Truhlar. Universal solvation model based on solute electron density and on a continuum model of the solvent defined by the bulk dielectric constant and atomic surface tensions. The Journal of Physical Chemistry B, 113(18):6378–6396, 2009. PMID: 19366259.

[14] A. Warshel, K. S. Pankaz, K. Mitsunori, and W. W. Parson. Modeling electrostatic effects in proteins. Biochimica et Biophysica Acta (BBA) - Proteins and Proteomics, 1764(11):1647–1676, 2006.

[15] R. C. Harris B., and B. M. Pettitt. Effects of geometry and chemistry on hydrophobic solvation. Proc. Natl. Acad. Sci. USA, 111(41):14681–14686, 2014.

[16] F. Eisenhaber. Hydrophobic regions on protein surfaces. derivation of the solvation energy from their area distribution in crystallographic protein structures. Protein Science, 5(8):1676–1686, 1996.

[17] Y. H. Singh, M. M. Gromiha, A. Sarai, and S. Ahmad. Atom-wise statistics and prediction of solvent accessibility in proteins. Biophysical Chemistry, 124(2):145 – 154, 2006.

[18] M. Stöhr and A. Tkatchenko. Quantum mechanics of proteins in explicit water: The role of plasmon-like solute-solvent interactions. Science Advances, 5(12), 2019.

[19] R. B. Hermann. Theory of hydrophobic bonding. II. correlation of hydrocarbon solubility in water with solvent cavity surface area. The Journal of Physical Chemistry, 76(19):2754–2759, 1972.

[20] F. M. Richards. Areas, volumes, packing, and protein structure. Annual Review of Biophysics and Bioengineering, 6(1):151–176, 1977.

[21] J. Greer and B. L. Bush. Macromolecular shape and surface maps by solvent exclusion. Proc. Natl. Acad. Sci. USA, 75(1):303–307, 1978.

[22] C. Chothia. The nature of the accessible and buried surfaces in proteins. Journal of Molecular Biology, 105(1):1–12, 1976.

[23] J. Janin. Surface and Inside Volumes in Globular Proteins. Nature, 277:491–492, February 1979.

[24] S. Miller, J. Janin, A. M. Lesk, and C. Chothia. Interior and surface of monomeric proteins. Journal of Molecular Biology, 196(3):641–656, 1987.

[25] G. J. Lesser and G. D. Rose. Hydrophobicity of amino acid subgroups in proteins. Proteins: Structure, Function, and Bioinformatics, 8(1):6–13, 1990.

[26] Y. Harpaz, M. Gerstein, and C. Chothia. Volume changes on protein folding. Structure, 2(7):641–649, 1994.

[27] R. L. Baldwin and G. D. Rose. How the hydrophobic factor drives protein folding. Proc. Natl. Acad. Sci. USA, 113(44):12462–12466, 2016.

[28] A. L. Lomize, I. D. Pogozheva, and H. I. Mosberg. Anisotropic solvent model of the lipid bilayer. 1. parameterization of long-range electrostatics and first solvation shell effects. Journal of Chemical Information and Modeling, 51(4):918–929, 2011. PMID: 21438609.

[29] A. T. Brunger. Version 1.2 of the Crystallography and NMR System. Nature Protocol, 2:2728–2733, February 2007.

[30] J. M. Word, S. C. Lovell, J. S. Richardson, and D. C. Richardson. Asparagine and glutamine: using hydrogen atom contacts in the choice of side-chain amide orientation. Journal of Molecular Biology, 285(4):1735–1747, 1999.

[31] B. R. Brooks, C. L. Brooks, A. D. Mackerell, L. Nilsson, R. J. Petrella, B. Roux, Y. Won, G. Archontis, C. Bartels, S. Boresch, A. Caflisch, L. Caves, Q. Cui, A. R. Dinner, M. Feig, S. Fischer, J. Gao, M. Hodoscek, W. Im, K. Kuczera, T. Lazaridis, J. Ma, V. Ovchinnikov, E. Paci, R. W. Pastor, C. B. Post, J. Z. Pu, M. Schaefer, B. Tidor, R. M. Venable, H. L. Woodcock, X. Wu, W. Yang, D. M. York, and M. Karplus. Charmm: The biomolecular simulation program. Journal of Computational Chemistry, 30(10):1545–1614, 2009.

[32] W. Kabsch and C. Sander. Dictionary of protein secondary structure: pattern recognition of hydrogen-bonded and geometrical features. Biopolymers, 22(12):2577– 2637, 1983.

[33] D. Eisenberg and A. D. McLachlan. Solvation energy in protein folding and binding. Nature, 319:199–203, 1986.

[34] M. Delarue and P. Koehl. Atomic environment energies in proteins defined from statistics of accessible and contact surface areas. Journal of Molecular Biology, 249(3):675–690, 1995.

[35] D. L. Mobley, A. E. Barber II, C. J. Fennell, and K. A. Dill. Charge asymmetries in hydration of polar solutes. The Journal of Physical Chemistry B, 112(8):2405– 2414, 2008. PMID: 18251538.

[36] L. Lins and R. Brasseur. The hydrophobic effect in protein folding. The FASEB Journal, 9(7):535–540, 1995.

[37] T. Simonson and D. Perahia. Internal and interfacial dielectric properties of cytochrome c from molecular dynamics in aqueous solution. Proc. Natl. Acad. Sci. USA, 92(4):1082–1086, 1995.

[38] L. Li, C. Li, Z. Zhang, and E. Alexov. On the dielectric “constant” of proteins: Smooth dielectric function for macromolecular modeling and its implementation in delphi. Journal of Chemical Theory and Computation, 9(4):2126–2136, 2013. PMID: 23585741.

[39] M. Gerstein and C. Chothia. Packing at the protein-water interface. Proc. Natl. Acad. Sci. USA, 93(19):10167–10172, 1996.

[40] D. Laage, T. Elsaesser, and J. T. Hynes. Water dynamics in the hydration shells of biomolecules. Chemical Reviews, 117(16):10694–10725, 2017.

[41] A. Pintar, O. Carugo, and S. Pongor. Atom depth as a descriptor of the protein interior. Biophysical Journal, 84(4):2553–2561, 2003.

[42] T. M. Raschke, J. Tsai, and M. Levitt. Quantification of the hydrophobic interaction by simulations of the aggregation of small hydrophobic solutes in water. Proc. Natl. Acad. Sci. USA, 98(11):5965–5969, 2001.

[43] L. Wang, Y. Pang, T. Holder, J. R. Brender, A. V. Kurochkin, and E. R. P. Zuiderweg. Functional dynamics in the active site of the ribonuclease binase. Proc. Natl. Acad. Sci. USA, 98(14):7684–7689, 2001.

[44] J. Huang, P. E. M. Lopes, B. Roux, and A. D. MacKerell. Recent advances in polarizable force fields for macromolecules: Microsecond simulations of proteins using the classical drude oscillator model. The Journal of Physical Chemistry Letters, 5(18):3144–3150, 2014. PMID: 25247054.

## References

[1] S. F. Altschul, T. L. Madden, A. A. Schäffer, J. Zhang, Zheng Zhang, W. Miller, and D. J. Lipman. Gapped BLAST and PSI-BLAST: a new generation of protein database search programs. Nucleic Acids Research, 25(17):3389–3402, 09 1997.

